# Unique spatial integration in mouse primary visual cortex and higher visual areas

**DOI:** 10.1101/643007

**Authors:** Kevin A. Murgas, Ashley M. Wilson, Valerie Michael, Lindsey L. Glickfeld

## Abstract

Neurons in the visual system integrate over a wide range of spatial scales. This diversity is thought to enable both local and global computations. To understand how spatial information is encoded across the mouse visual system, we use two-photon imaging to measure receptive fields in primary visual cortex (V1) and three downstream higher visual areas (HVAs): LM (lateromedial), AL (anterolateral) and PM (posteromedial). We find significantly larger receptive field sizes and less surround suppression in PM than in V1 or the other HVAs. Unlike other visual features studied in this system, specialization of spatial integration in PM cannot be explained by specific projections from V1 to the HVAs. Instead, our data suggests that distinct connectivity within PM may support the area’s unique ability to encode global features of the visual scene, whereas V1, LM and AL may be more specialized for processing local features.

## Introduction

Hierarchical and parallel processing are two major organizing principles of sensory systems (Ungerleider and Mishkin, 1982; Felleman and Van Essen, 1991; Goodale and Milner, 1992; Nassi and Callaway, 2009). Together, these principles support an increase in both the specialization and generalization of receptive fields along feed-forward sensory processing pathways (Zeki, 1978; Kobatake and Tanaka, 1994). Specialization within an area both 1) enables better discrimination of selected features and 2) allows for a distinct set of computations to be performed across areas. In comparison, generalization is thought to allow for invariant representations of selected features amid variation in other features (e.g. position, size, or viewing angle (Riesenhuber and Poggio, 2002)). Thus, the transformations that occur along the visual hierarchy mediate increases in both selectivity and tolerance of receptive fields to support higher-level perception (Rust et al., 2006; DiCarlo et al., 2012).

Our ultimate goal is to determine the circuit mechanisms that support such transformations of receptive fields across visual areas. The mouse is a useful model for studying such mechanisms, and as in humans and non-human primates, mice have an array of higher visual areas (HVAs) that each form their own, nearly complete, representation of the visual field (Wang and Burkhalter, 2007; Garrett et al., 2014; Glickfeld and Olsen, 2017). Also like primates, the functional properties of the neurons in the HVAs are more specialized for selected features than the population of neurons in V1 (Andermann et al., 2011; Marshel et al., 2011; Roth et al., 2012). Target-specific anatomical connectivity between V1 and the HVAs underlies at least some of this increase in specialization (Glickfeld et al., 2013; Matsui and Ohki, 2013; Han et al., 2018). However, as of yet, very little evidence for increasing generalization has been identified in the HVAs of the mouse (Juavinett and Callaway, 2015). Thus, in order for the mouse to be a useful model for understanding the diversity of mechanisms that underlie the transformations of receptive fields, we need a more complete understanding of the types of transformations that occur.

The scale of spatial integration is an important determinant of higher-order representations in the visual system. Receptive field size typically increases along the visual hierarchy (Dräger, 1975; Wang and Burkhalter, 2007; Vermaercke et al., 2014) and is thought to be necessary for the generation of position- and size-invariant responses (Rust and DiCarlo, 2010; Tafazoli et al., 2017). Yet, the responses of neurons to stimuli of different sizes is not simply defined by their receptive field size. Interactions in the extra-classical receptive field often drive suppression of responses to larger stimuli and generate size tuning (Hubel and Wiesel, 1968; DeAngelis et al., 1994; Angelucci et al., 2017). Suppression of responses by larger stimuli may support the generation of receptive fields specialized for local computations such as identification of boundaries or objects (Lamme et al., 1995; Coen-Cagli et al., 2012). Conversely, integration across large spatial scales might enable generalization of feature representation across sizes. In addition, the lack of surround suppression in the dorsal stream of non-human primates has been proposed to support specialization of these areas for encoding global motion and optic flow (Tanaka et al., 1986; Born and Tootell, 1992). Thus, investigating how size is encoded across the HVAs may reveal transformations that could support both generalization and specialization of receptive fields in the visual system.

To investigate how size is encoded in the mouse visual system, we used two-photon calcium imaging to measure receptive field size and size tuning in populations of layer 2/3 neurons in V1 and three HVAs that receive the strongest direct input from V1: LM (lateromedial), AL (anterolateral) and PM (posteromedial). Similar to previous observations, we find that receptive field sizes in the HVAs are larger than in V1, and larger in PM than in AL or LM. Neurons in PM also had larger preferred sizes and much less surround suppression than was observed in the other three areas. These differences in preferred size and degree of suppression were conserved across stimulus contrasts and could not be explained by target-specific projections from V1. This suggests that there may be fundamental differences in connectivity and the recruitment of normalization mechanisms across the HVAs. Moreover, the unique encoding of size in PM argue that it is poised to transform signals into increasingly general representations of the external world.

## Results

### Visual Cortical Areas Have Distinct Receptive Field Sizes

To examine the representation of size in V1 and the HVAs, we first characterized the spatial receptive fields of neurons in these areas. Using widefield calcium imaging in awake mice transgenically expressing the calcium indicator GCaMP6 in pyramidal cells (GCaMP6f: n=7 and GCaMP6s: n=2), we generated retinotopic maps of visual cortex (Figure 1A-B). V1 and the HVAs were designated based on previously established HVA maps (Wang and Burkhalter, 2007; Andermann et al., 2011). We then used these maps to target our two-photon (2P) imaging experiments to the area of interest, as well as to inform the approximate position of spatial receptive fields for those neurons.

**Figure 1.**
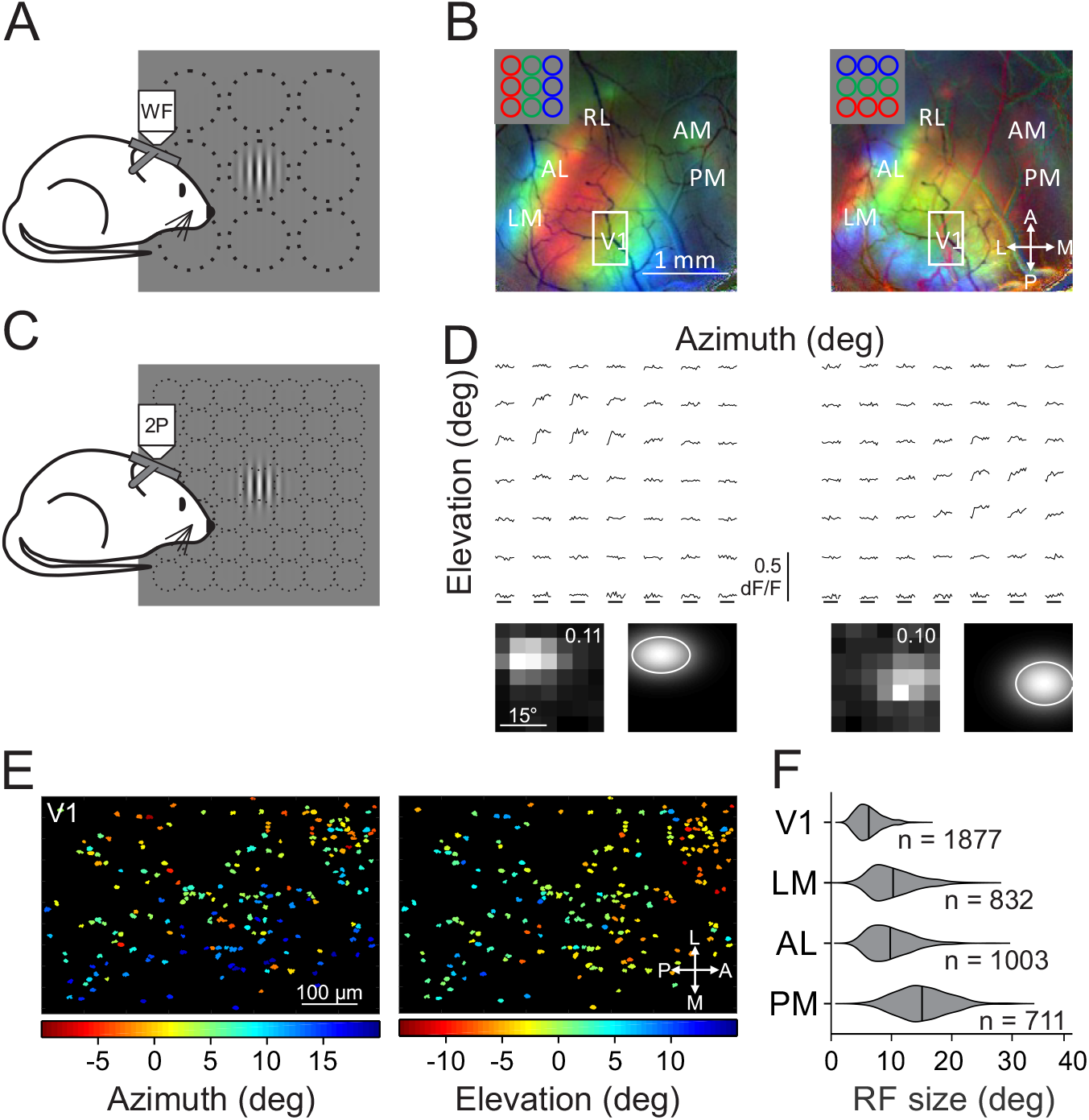
Retinotopic mapping of V1 and HVAs reveals differences in receptive field size across areas. **A.** Schematic of rough retinotopy experiments. Under widefield fluorescence microscopy, mice were presented 30º diameter Gabor gratings at one of 9 positions in a 3×3 grid. **B.** Pseudocolored retinotopic maps showing changes in fluorescence (dF/F) in response to stimuli across three azimuths (left) or elevations (right) for an example mouse. **C.** Schematic of fine retinotopy experiments. Under 2-photon fluorescent microscopy, mice were presented Gabor gratings at one of 49 positions in a 7×7 grid. **D.** Top, average dF/F time-courses for two example cells in response to stimuli in each of the 49 positions. Bottom, average responses (left; value is the maximum dF/F) and 2-dimensional gaussian fits (right; contour is 1 sigma) for the cells above. **E.** Receptive field (RF) center azimuth (left) and elevation (right) for all cells in the field of view. Field of view from white rectangle in **B**. **F.** Summary of RF size (half-width at half-max (HWHM)) for all neurons in each area (n=9 mice). All comparisons, except LM-AL (p=0.14), are significantly different (p<0.001).

Using cellular-resolution two-photon (2P) imaging, we examined the spatial receptive field organization of pyramidal cells in layer 2/3 (L2/3) of V1 and the three largest and best-characterized HVAs (LM, AL, PM). To measure receptive fields, we presented small (5-10°; see Methods) drifting gratings in a 7×7 grid (Figure 1C-D). Averaged responses to the 49 stimulus positions were fit with a 2-dimensional Gaussian function to model the cell’s receptive field size and position in retinotopic space (Figure 1D-E). These experiments revealed differences in receptive fields across V1 and the HVAs. Consistent with previous studies (Wang and Burkhalter, 2007), receptive field sizes in V1 were significantly smaller than in all of the other areas (half-width at half-max (HWHM): 6.2°±0.1° (mean ± STD), n=1877 cells, 9 mice; Kruskal-Wallis test (p<0.001) with post-hoc Tukey tests all comparisons with V1: p<0.001; Figure 1F). Of the HVAs, LM and AL exhibited intermediate receptive field sizes which were not significantly different from each other (LM: 10.3°±0.1°, n=832 cells, 9 mice; AL: 9.8°±0.1°, n=1003 cells, 9 mice; post-hoc Tukey test LM-AL: p=0.14). Finally, the receptive fields of neurons in PM were significantly larger than those in both AL and LM (15.1°±0.1°, n=711 cells, 9 mice; post-hoc Tukey tests all comparisons with PM: p<0.001). Thus, we find a general increase in receptive field size from L2/3 of V1 to L2/3 of the HVAs, consistent with an increase in generalization in the HVAs. Further, we find a difference across areas, consistent with a specialization of the function of the HVAs.

### Visual Cortical Areas Have Distinct Preferred Sizes and Surround Suppression

To further investigate spatial integration in V1 and the HVAs, we measured single neuron responses to stimuli of varying diameter and contrast (Figure 2A). We first addressed size tuning of neurons at the highest contrast used (0.8). Since size tuning is sensitive to how well the stimulus is centered on the receptive field, we only examined neurons with well-fit receptive fields close to the center of the size-tuning stimulus (see Methods). Size-tuning curves lie along a continuum which can be described by either a single sigmoid (SS), where responses are monotonically increasing or saturating, or a difference of sigmoids (DOS), in which responses initially increase but then are suppressed at larger sizes (Sceniak et al., 1999). Thus, we fit all cells with both models and used a sequential F-test to statistically determine if the DOS model was significantly better than the SS model (Figure 2B-C). This analysis revealed significant differences in the size tuning of neurons across areas (two-way ANOVA: main effect of area: p<0.001, interaction of area and size: p<0.001; Figure 2D).

**Figure 2.**
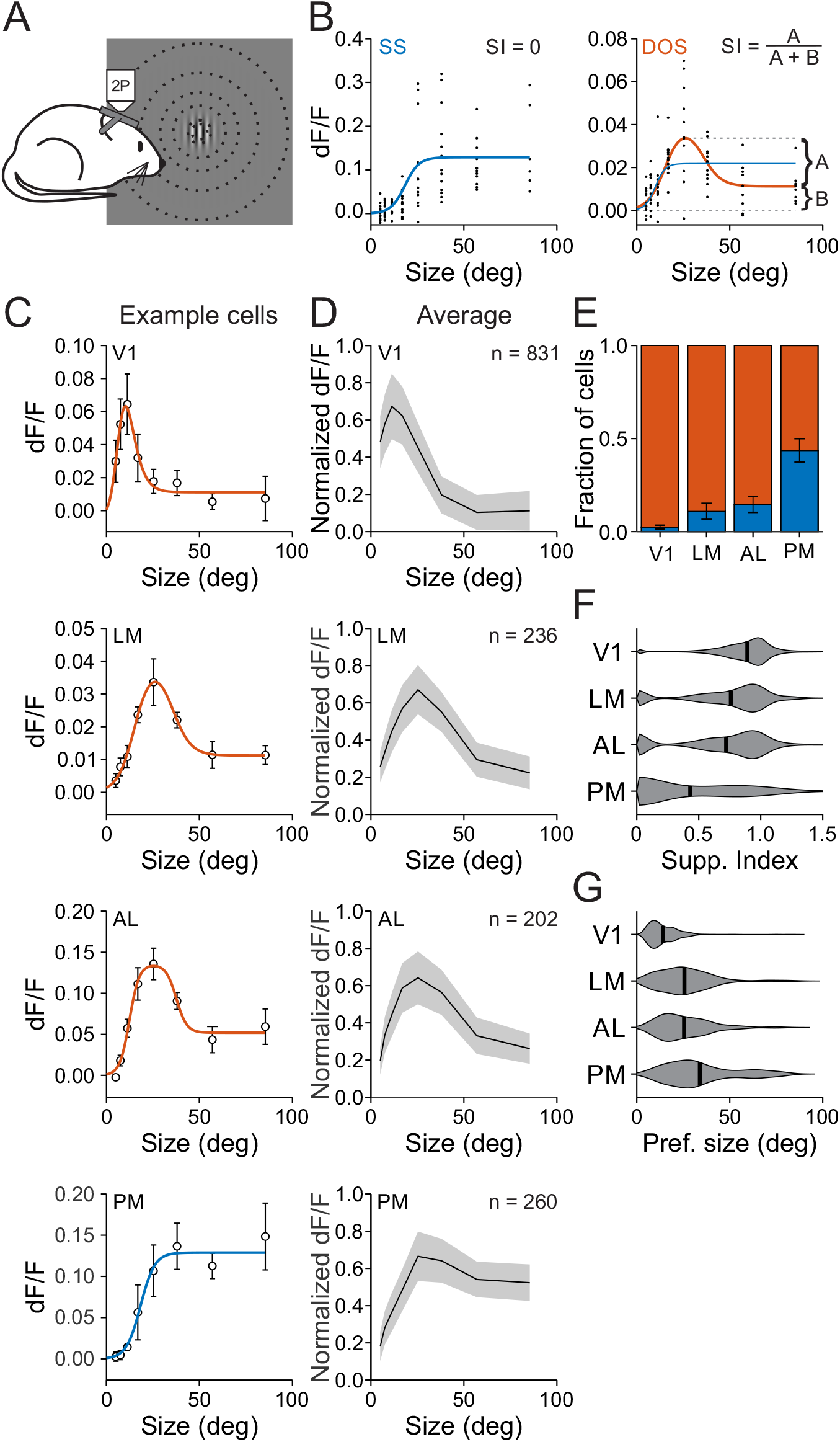
Size is encoded differently across visual cortical areas. **A.** Schematic of size-tuning experiments. Under 2P fluorescent microscopy, mice are presented Gabor gratings of varying size at a fixed position. **B.** Single trial responses (black dots) to stimuli of varying size fit with either a single sigmoid (SS, left, blue) or both a single sigmoid and a difference of sigmoids (DOS, right, red) for two example cells. If the SS is the better fit then, the suppression index (SI) is zero; if the DOS is the better fit, then the SI is equal to the ratio of the dF/F at the preferred size and the difference between the dF/F at the preferred size and the largest size. **C.** Average dF/F responses and best fit for an example cell in each area. Cells in LM and PM are the cells in **B**. Error is ± SEM across trials. **D.** Average responses to each size stimulus across all well-fit cells in each area. Error is ± STD across cells. **E.** Proportion of cells best fit by an SS (blue) or DOS (red). Error is ± SEP across cells. p<0.001, Chi-squared test. **F.** Summary of suppression index by area. All comparisons, except LM-AL (p=0.79), are significantly different (p<0.001). Violins are kernel density estimators; thick bars are mean. **G.** Summary of preferred size by area. All comparisons, except LM-AL (p=1.00), are significantly different (p<0.001).

Differences in size tuning across visual areas were related to a number of tuning features of the population. First, the prevalence of suppressed cells was significantly different across areas. The highest fraction of suppressed cells was in V1, the lowest was in PM, and there was an intermediate fraction in LM and AL (Fraction suppressed cells-V1: 0.976±0.005 (mean±SEP), n=831 cells; LM: 0.891±0.022, n=202 cells; AL: 0.854±0.022, n=260 cells; PM: 0.564±0.032, n=236 cells; Chi-squared test: p<0.001; Figure 2E). Second, the amount of suppression, measured as the suppression index (SI), was significantly different across areas. Neurons in V1 had the highest SI, PM had the lowest SI, and LM and AL were not statistically different from each other (SI-V1: 0.89±0.01; LM: 0.76±0.02; AL: 0.72±0.02; PM: 0.43±0.02; Kruskal-Wallis test (p<0.001) with post-hoc Tukey tests: all comparisons except LM-AL: p<0.001; LM-AL: p=0.79; Figure 2F). This difference could not be explained by the area-specific inclusion criteria based on the distance between the receptive field and stimulus center. If we used the same cutoff in all areas (15º), the significant differences observed above remained present (SI with matched cutoff - V1: 0.86±0.01, n=1092 cells; LM: 0.76±0.02, n=202; AL: 0.72±0.02, n=260; PM: 0.49±0.03, n=169; Kruskal-Wallis test (p<0.001) with post-hoc Tukey tests: all comparisons except LM-AL: p<0.005; LM-AL: p=0.79). Moreover, this difference in SI could not be explained by the different fractions of suppressed and non-suppressed neurons since the difference across areas held even when we only considered suppressed cells (SI in suppressed cells - V1: 0.91±0.01, n=811 cells; LM: 0.85±0.02, n=180; AL: 0.84±0.02, n=222; PM: 0.77±0.02, n=133; Kruskal-Wallis test (p<0.001) with post-hoc Tukey tests: all comparisons with V1: p<0.005; LM-AL: p=0.98; LM-PM: p<0.05; AL-PM: p=0.07).

Finally, there were significant differences across areas in the preferred size of neurons. The preferred size of neurons in V1 was significantly smaller than those in the HVAs (preferred size-V1: 14.1°±0.4°; Kruskal-Wallis test (p<0.001) with post-hoc Tukey tests: all comparisons with V1: p<0.001; Figure 2G). Within HVAs, the preferred size of neurons in LM and AL were not significantly different from each other, but were both significantly smaller than the neurons in PM (LM: 25.6°±0.9°; AL: 25.5°±0.8°; PM: 33.9°±0.8°; post-hoc Tukey test: all comparisons with PM: p<0.001; LM-AL: p=1.00). Notably, PM was the only area where there was a significant correlation between the receptive field size and preferred size within individual cells, consistent with the receptive field being a stronger determinant of spatial integration in PM (slope=0.36; p<0.01; linear regression). Thus, these data suggest that PM has a unique representation of stimulus size compared to V1 and the other HVAs, and therefore may be specialized for processing global features of the visual scene.

### Contrast sensitivity of neurons is similar across areas

There is a strong relationship between surround suppression and stimulus contrast (Sceniak et al., 1999). Thus, we next explored whether the differences in size tuning that we observed across areas could be explained by differences in contrast sensitivity. We identified cells that were well-fit for size, and extrapolated contrast response functions from the fits at the preferred size at the highest contrast (Figure 3A-B). We then fit these data with the Naka-Rushton function to measure the contrast at 50% response (C_50_). While the average contrast response functions across areas were similar, there were significant differences across areas (two-way ANOVA: main effect of area: p<0.001; interaction of area and contrast: p<0.05; Figure 3C). This was largely due to neurons in AL having a higher C_50_ than the other areas, although this was only significant in comparison to V1 (V1: 0.34±0.01, n=482 cells; LM: 0.34±0.02, n=103; AL: 0.39±0.01, n=168; PM: 0.34±0.01, n=131; Kruskal-Wallis test (p<0.01) with post-hoc Tukey tests: V1-LM: p=1.00; V1-AL: p<0.005; V1-PM: p=0.99; LM-AL: p=0.07; LM-PM: p=0.99; AL-PM: p=0.09; Figure 3D). However, when measured at the same size (20º) across neurons, instead of at each cell’s preferred size, there was no significant difference in the contrast response functions (two-way ANOVA: main effect of area: p=0.411; n=598 cells) or C_50_ (Kruskal-Wallis test: p=0.484) across areas. Thus, we conclude that if there are differences in contrast sensitivity across areas, they are weak, and potentially dependent on the differences in size tuning across areas.

**Figure 3.**
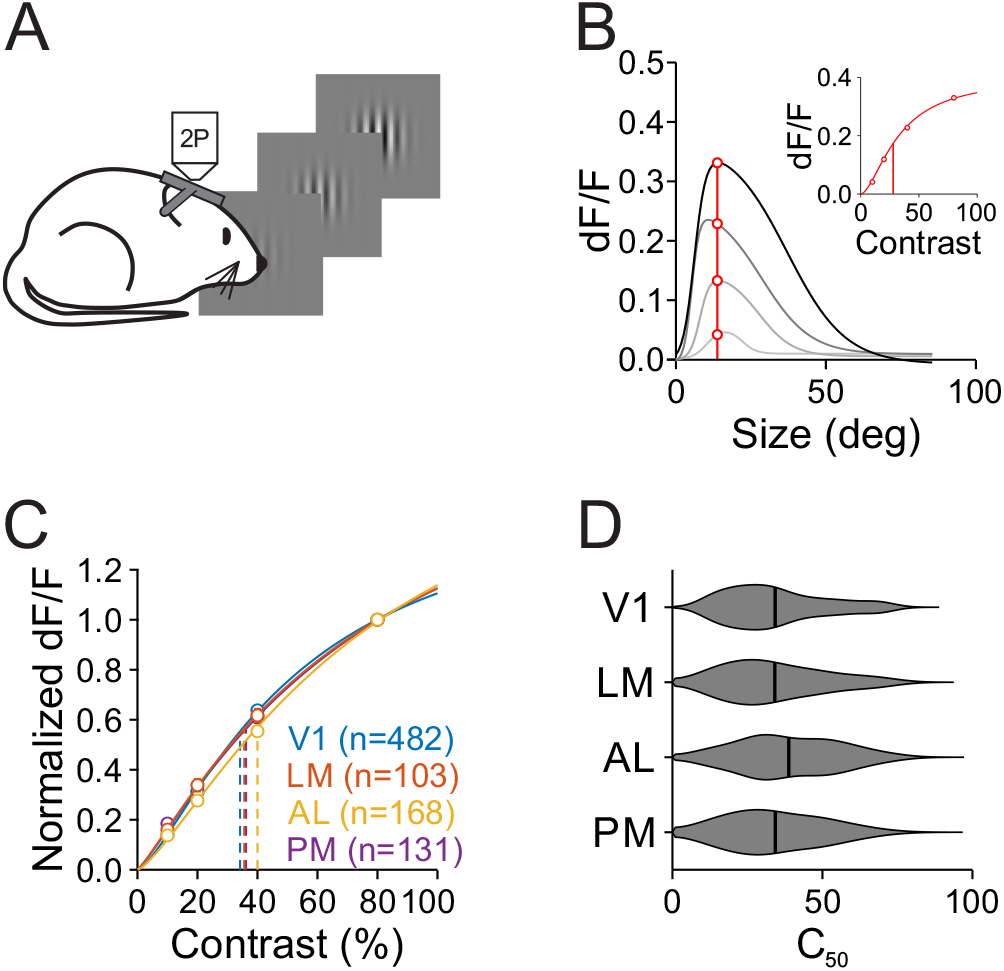
Contrast sensitivity is similar across areas. **A.** Schematic of size-by-contrast-tuning experiments. Under 2P fluorescent microscopy, mice are presented Gabor gratings of variable size and contrast at a fixed position. **B.** Contrast-response extraction in an example cell. Response values at the cell’s preferred size (red line and circles, found at the highest contrast) were extracted from size-tuning curves across four contrasts (gray curves). Inset-contrast response function for this cell fit with the Naka-Rushton equation. Vertical line is C_50_. **C.** Summary of averaged normalized contrast response functions across area. Error is ± SEM across cells. Fits are of average contrast response measures. Interaction of area and contrast: p<0.05. **D.** Summary of C_50_ for all cells in each area. Only V1 and AL are significantly different (p<0.005).

In addition, we find only weak dependence of size-tuning on stimulus contrast. In each area, we observed a strong effect of contrast on visual responses (two-way ANOVA: main effect of contrast for all areas: p<0.001; V1: n=233 cells; LM: n=87; AL: n=81; PM: n=113; Figure 4A). However, there was a similar proportion of suppressed cells at high and low contrast (0.1 vs 0.8 – Chi-squared tests: V1: p=0.43; LM: p=0.19; AL: p=0.06; PM: p=0.42; Figure 4B). While there was no significant contrast-dependence of SI (two-way ANOVA: main effect of contrast: p=0.36; interaction of contrast and area: p=0.55; Figure 4C), preferred size was significant contrast-dependent within each area (two-way ANOVA: main effect of contrast: p<0.05; interaction of contrast and area: p=0.81; Figure 4D). Importantly, however, the lowest-contrast data still reflected significant differences in size tuning across areas (Fraction suppressed: Chi-squared test: p<0.001; SI: Kruskal-Wallis test (p<0.001) with post-hoc Tukey tests all comparisons except LM-AL: p<0.001; LM-AL: p=0.36; preferred size: Kruskal-Wallis test (p<0.001) with post-hoc Tukey tests: all comparisons except LM-AL and AL-PM: p<0.001; LM-AL: p=0.17; AL-PM: p=0.17). Thus, neurons in V1 have narrower spatial integration and neurons in PM have broader integration, independent of stimulus contrast.

**Figure 4.**
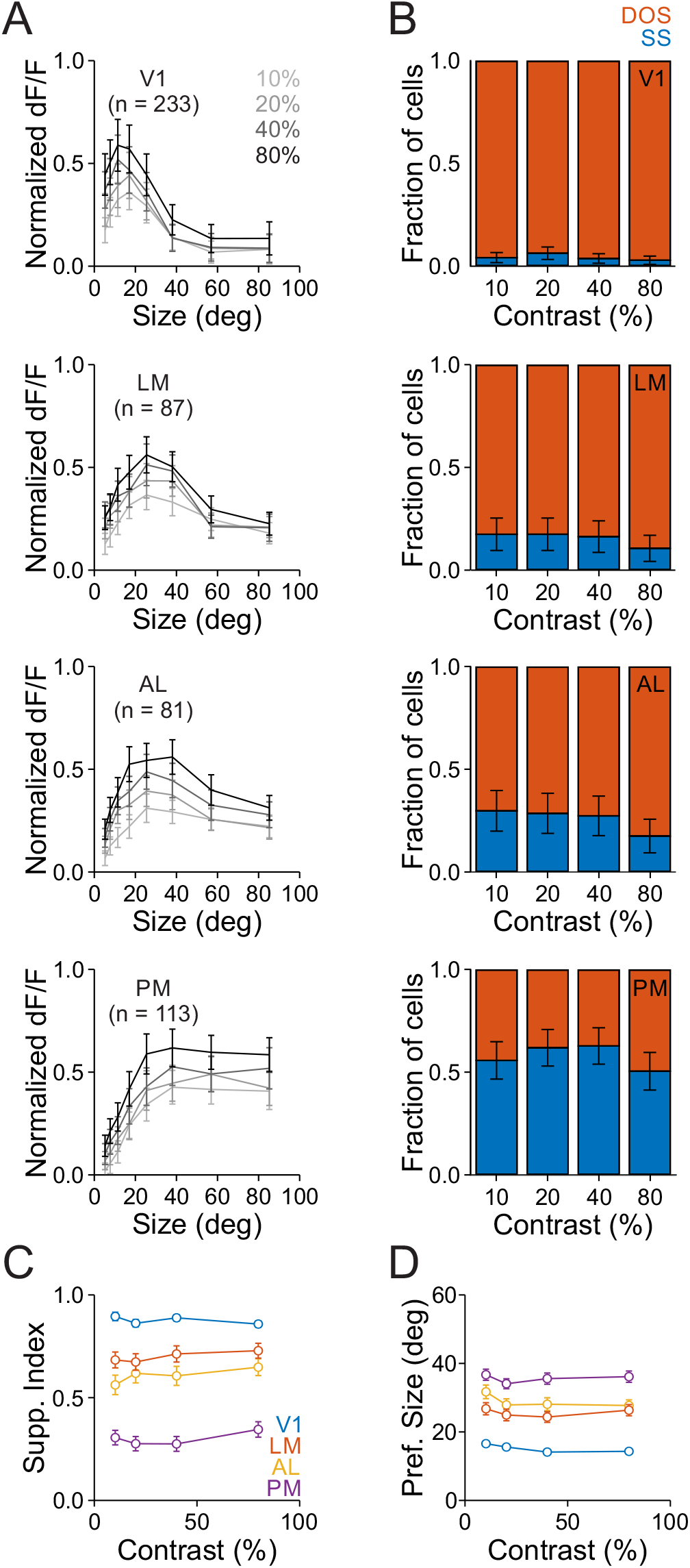
Differences in size tuning in V1 and the HVAs is conserved across contrasts. **A.** Average responses of all cells in each area to each size for all contrasts. Error is ± SEM across cells. **B.** Proportion of cells best fit by an SS (blue) or DOS (red) for each contrast in each area. Error is ± SEP across cells. 10% vs 80% contrast p>0.05 for all areas. **C.** Summary of suppression index as a function of contrast in each area. Error is ± SEM across cells. No main effect of contrast: p=0.36. **D.** Same as **C**, for preferred size. Main effect of contrast: p<0.05.

### Differences in spatial integration in the HVAs are not inherited from V1

Our experiments have revealed that neurons in the HVAs integrate over a much larger visual field than neurons in V1. In addition, we find that there is specialization among the HVAs where PM integrates more broadly than AL and LM. To test whether these difference can be explained by functionally specific projections to the HVAs, we virally expressed GCaMP6s in V1 neurons (Figure 5A-B), and used 2P imaging to measure the functional properties of V1 inputs to LM, AL and PM (Figure 5C; (Glickfeld et al., 2013)). For these experiments, we used a similar set of stimuli as when imaging cell bodies: first, using a dense presentation of stimuli in 49 different positions to measure receptive field organization (Figure 5D-F); and second, varying stimulus size (at a contrast of 0.8) to measure the preferred size and SI (Figure 6).

**Figure 5.**
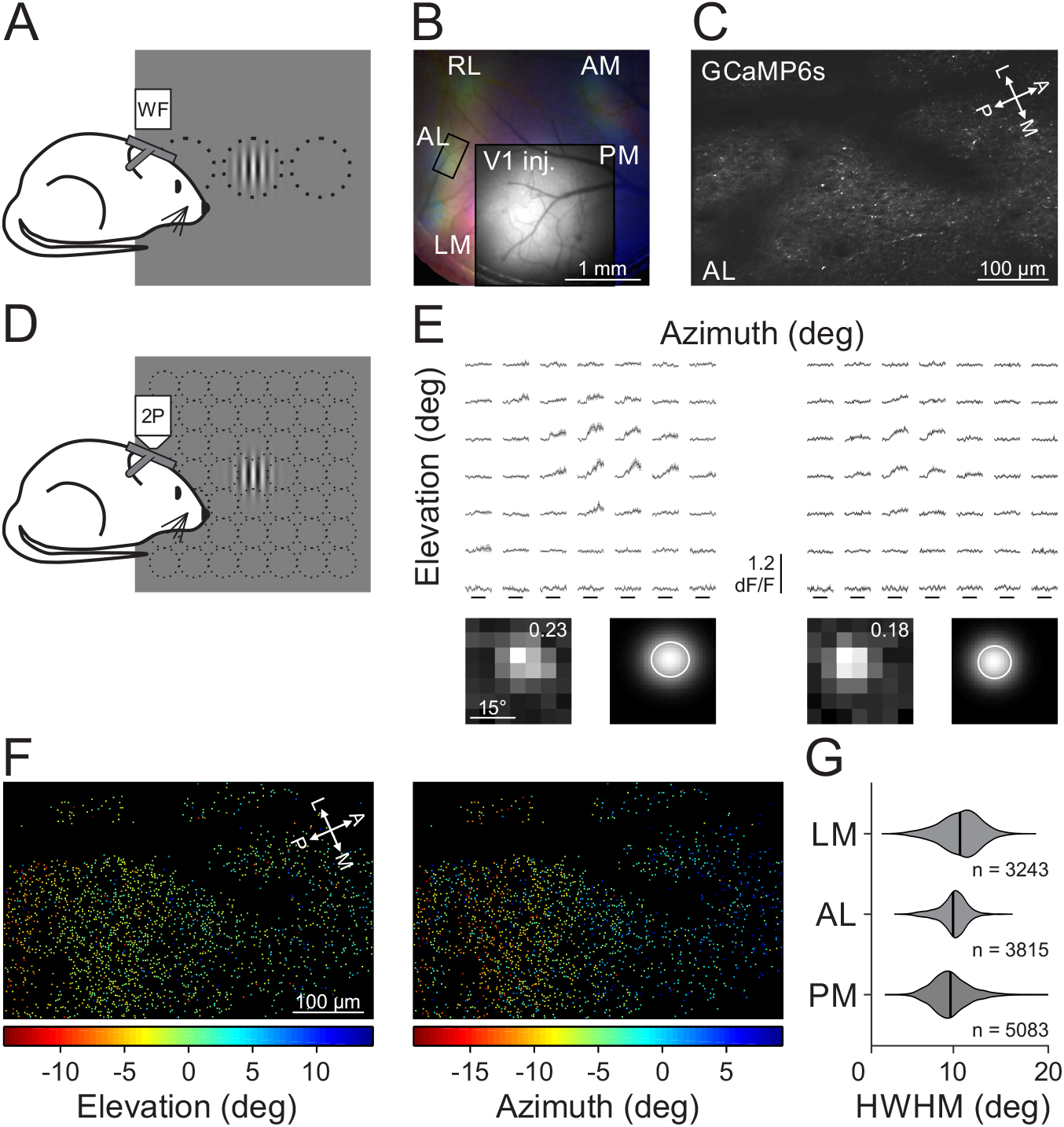
Receptive field size of V1 inputs to the HVAs is similar across areas. **A.** Schematic of widefield imaging stimulus and setup. **B.** Pseudo-color image of changes in fluorescence (dF/F) in axonal projections from V1 to the HVAs in response to stimuli of different azimuth (same conventions as Figure 1B) for an example mouse. Inset: Raw fluorescence (F) image of the injection site in V1. **C.** Average fluorescence image of an example field of view from the region of interest in **B** (rectangle in AL). **D.** Schematic of fine retinotopy stimulus for 2P imaging. **E.** Average fluorescence (dF/F) traces (top) and RF with fit (bottom, same conventions as Figure 1D) for two example cells from the field of view in **C**. **F.** RF center azimuth (left) and elevation (right) for all V1 boutons in the field of view in **C**. **G.** Summary of RF size for V1 boutons in each higher visual area. All comparisons: p<0.001.

**Figure 6.**
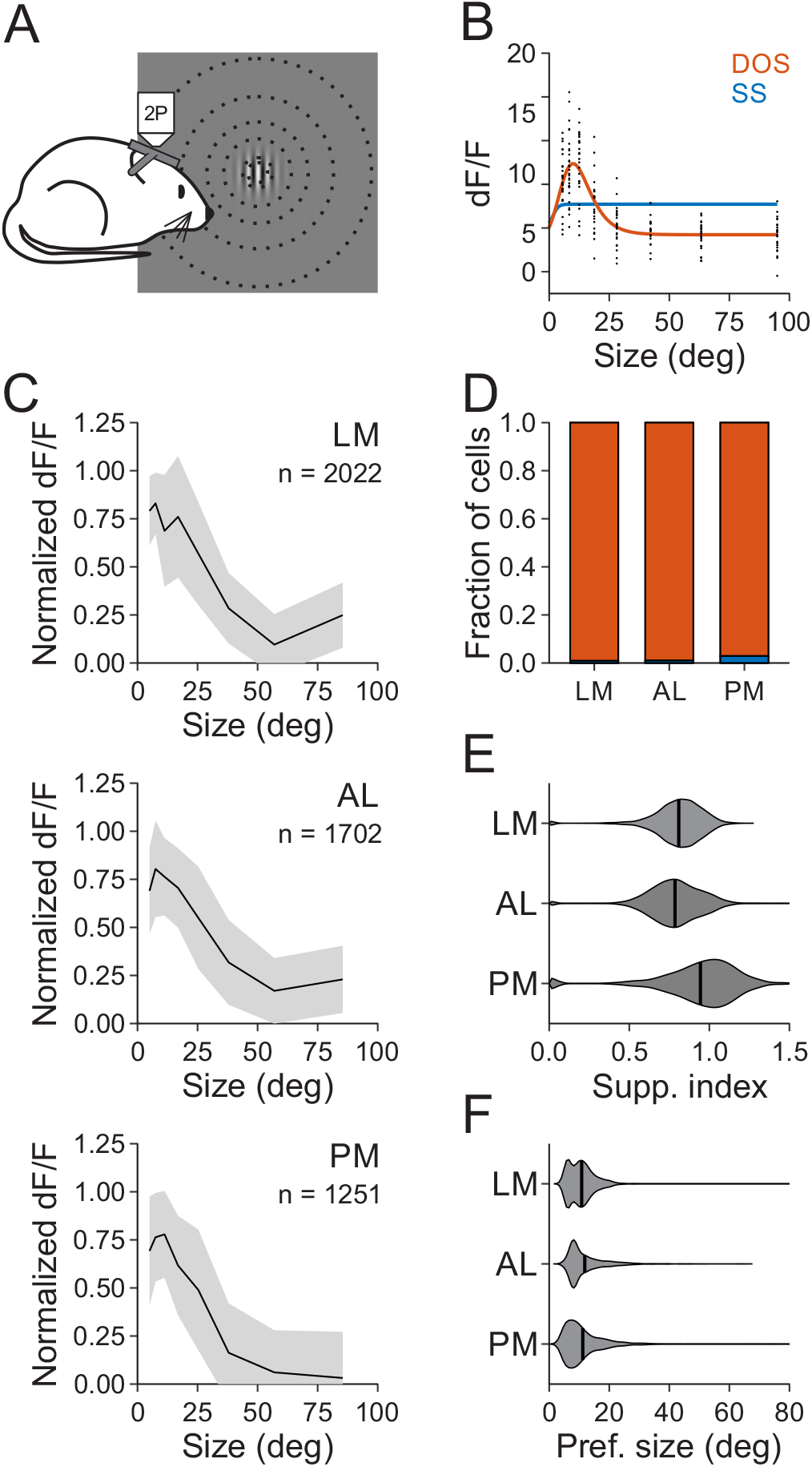
Size tuning of V1 inputs to the HVAs is similar across areas. **A.** Schematic of stimulus protocol for size tuning experiments. **B.** Example single trial (black) responses to stimuli of varying size for a single cell. Data is fit with either a SS (blue) or a DOS (red). **C.** Average normalized dF/F for all well-fit boutons in each area. Error is ± STD across cells. **D.** Proportion of boutons best fit by an SS (blue) or DOS (red) for each contrast in each area. Error is ± SEP across cells. Chi-squared test: p<0.001. **E.** Summary of suppression index for all well-fit boutons in each area. All comparisons: p<0.001. **F.** Same as **E**, for preferred size. All comparisons: p<0.001.

Receptive fields of V1 axons in the HVAs were similarly sized across areas (HWHM - LM: 10.0°±2.3°, n=3243 boutons, 4 mice; AL: 9.2°±1.6°, n=3815 boutons, 3 mice; PM: 8.9°±2.1°, n=5083 boutons, 4 mice; Kruskal-Wallis test: p<0.001; with post-hoc Tukey tests, all comparisons: p<0.001; Figure 5G). The receptive field size of V1 axons in the HVAs was significantly larger than the receptive field size of neurons in L2/3 of V1 (Kruskal-Wallis test (p<0.001) with post-hoc Tukey tests all comparisons with V1: p<0.001), perhaps due to the inclusion of axons from deeper layers in V1 where neurons have larger receptive fields (Niell and Stryker, 2008)). In fact, only in PM did the imaged neurons have significantly larger receptive fields than the V1 boutons coming into that area (Wilcoxon rank-sum test: p<0.001). Together, this suggests that there is relatively little specificity for receptive field size in the projections to the HVAs. Moreover, the increase in receptive field size between L2/3 of V1 and LM and AL may partly be explained by computations within V1, while the increase in receptive field size in PM requires the convergence of multiple V1 axons or local computations within PM.

To address whether the differences in surround suppression in the HVAs could be explained by specific projections to the HVAs, we fit SS and DOS models to the responses of axons in the HVAs to stimuli of increasing size (Figure 6A-C). Projections from V1 to the HVAs had similar, although significantly different, fractions of suppressed boutons (LM: 0.991±0.002 (mean±SEP), n=2002 boutons; AL: 0.990±0.002, n=1702 boutons; PM: 0.971±0.005, n=1251 boutons; Chi-squared test: p<0.001; Figure 6D), degree of surround suppression (SI - LM: 0.81±0.15; AL: 0.79±0.17; PM: 0.94±0.25; Kruskal-Wallis test: p<0.001; with post-hoc Tukey tests, all comparisons: p<0.001; Figure 6E), and preferred size (LM: 10.8°±5.5°; AL: 11.8°±6.5°; PM: 11.2°±7.4°; Kruskal-Wallis test: p<0.001; with post-hoc Tukey tests: LM-AL: p<0.05; LM-PM: p<0.05; AL-PM: p<0.001; Figure 6F). The fraction of suppressed boutons was similar to, though significantly greater than, that of the cell bodies imaged in V1 (Chi-squared test: p<0.001), and significantly higher than the cell bodies imaged in their respective areas (Wilcoxon rank-sum tests for all areas: p<0.001). Moreover, the degree of suppression was significantly greater in V1 boutons than in the cell bodies in their target areas (Kruskal-Wallis test: p<0.001; with post-hoc Tukey tests for all areas: p<0.001). Thus, we find that the inputs to all areas undergo a similar degree of surround suppression, and less surround suppression than the neurons in the target areas, suggesting that there are unique mechanisms of spatial integration within the HVAs, and especially in PM.

## Discussion

Like primates, mice have multiple higher-order visual cortical areas (HVAs) that likely serve specialized roles in visual processing (Glickfeld and Olsen, 2017). However, as yet, little is known about the specific features encoded in each area, and therefore what function each area might serve. Here, we used two-photon calcium imaging to determine how stimulus size and contrast are encoded in neurons in V1 and three HVAs: LM, AL and PM.

We observed a number of differences in the receptive field properties of neurons in V1 and the HVAs. For instance, we found that receptive field size was significantly larger in the HVAs than in V1, and larger in PM than in AL or LM. This is consistent with electrophysiological measurements made from these areas (Dräger, 1975; Wang and Burkhalter, 2007), and the increase in receptive field size along the visual hierarchy of non-human primates (Baker et al., 1981). The increase in receptive field size may be a consequence of the increase in cortical magnification in the HVAs (Garrett et al., 2014), requiring increased convergence. Indeed, the inputs from V1 to each of the HVAs were very similarly sized, suggesting that the differences between PM and the other HVAs are not directly inherited from V1, as was found to be the case for other stimulus features (Glickfeld et al., 2013; Matsui and Ohki, 2013). Instead, these data suggest that there must be differences in convergence, local recurrent circuitry or the degree of feedback from other areas (e.g. other higher order cortical or thalamic areas).

Consistent with differences in local and feedback circuits, we also observed major differences in how these areas encode size information. Neurons in V1 were much more likely to be suppressed by large stimuli than neurons in any of the three HVAs. Among HVAs, PM had significantly fewer suppressed cells than either LM or AL. Even at the highest contrast, nearly 50% of cells in PM were better fit by a model that had no suppression at large sizes. Interestingly, there were very few non-suppressed inputs from V1 to any of the HVAs. In fact, we found a stronger suppression of V1 inputs to PM than to LM or AL. This suggests that the differences in encoding of size cannot be inherited from V1 either through specific inputs. If the spatial receptive field size reflects the degree of spatial integration in PM, then this also limits the degree to which convergence of inputs from V1 could support responses at the largest sizes. Instead, the transformation through which PM becomes sensitive to larger stimuli must be mediated by differences in local or feedback circuits.

In V1, somatostatin expressing interneurons are thought to pool excitatory inputs over long ranges and thereby mediate surround suppression (Adesnik et al., 2012; Dipoppa et al., 2018). However, anatomical data suggests that there are actually fewer somatostatin cells in V1 than in PM, and similar densities in PM, LM and AL (Kim et al., 2017). This suggests that there must be other differences in the structure of local connectivity, or the cell-types that support surround suppression, within these areas. Future experiments will be necessary to reveal whether surround suppression is the only form of normalization reduced in PM, or whether other forms of normalization, such as cross-orientation suppression, are also less pronounced.

In other species, surround suppression has been observed to persist or even increase along the visual hierarchy (Hubel and Wiesel, 1965; Schein and Desimone, 1990). Surround suppression has been suggested to be important for mediating computations involving salience, pop-out, or figure-ground segregation (Coen-Cagli et al., 2012). While AL and LM undergo less surround suppression than V1, there is still substantial suppression in these areas. This is consistent with the proposed participation of LM in the ventral visual stream and object recognition (Wang et al., 2011) and may support a role for AL in local motion signals. Meanwhile, populations of cells that lack surround suppression have been observed in both MT and MST (Tanaka et al., 1986; Born and Tootell, 1992). These widefield responses are proposed to support encoding of optic flow or other global motion signals. Lack of surround suppression may also enable selective responses to looming stimuli. Notably, similar to what has been observed in MT, only around half of neurons in PM lack surround suppression (Born and Tootell, 1992). This could enable parallel processing of both local and global computations in PM through functionally segregated networks.

Our data reveal a novel dimension for specialization of function in the HVAs: PM appears to be unique among the three HVAs examined in its representation of size. The larger receptive fields and lack of surround suppression in PM may support an increase in generalization by allowing for position invariance of visual response and integration of global motion signals across the larger regions of visual scenes. We have shown that this specialization does not arise from specific connectivity between V1 and the HVAs. Instead, we propose that it is due to unique connectivity among local circuits and cell-types in PM. Revealing the circuit mechanisms underlying the decrease in surround suppression will not only support our understanding of hierarchical transformations but also the mechanisms of cortical normalization. Thus, this work supports the use of the mouse as a model for studying higher visual cortical function.

## Methods

### Animals

All animal procedures conformed to standards set forth by the NIH, and were approved by the IACUC at Duke University. 14 mice (both sexes; 2-12 months old; singly and group housed (1-4 in a cage) under a regular 12-h light/dark cycle; C57/B6J (Jackson Labs #000664) was the primary background with up to 50% CBA/CaJ (Jackson Labs #000654)) were used in this study. Ai93 (*tm93.1(tetO-GCaMP6f)Hze*, Jackson Labs #024103; n = 2) and Ai94 (*tm94.1(tetO-GCaMP6s)Hze*, Jackson Labs #024104; n = 7) were crossed to *EMX1-IRES-Cre* (Jackson Labs #005628) and *CaMK2a-tTA* (Jackson Labs #003010) to enable constitutive GCaMP6 expression for cell body imaging. In order to decrease the likelihood of seizures (Steinmetz et al., 2017), transgenic mice were fed a diet of Doxycycline chow (200 mg/mL) (from onset of pregnancy until postnatal day 45 (P45)) to suppress calcium indicator expression.

### Cranial Window Implant and Viral Injection Surgeries

Cranial window surgeries were performed on mice older than P45 (Goldey et al., 2014). Animals were administered dexamethasone (3.2 mg/kg, s.c.) and meloxicam (2.5 mg/kg, s.c.) 2-6 hours prior to surgery, and anesthetized with ketamine (200 mg/kg, i.p.), xylazine (30 mg/kg, i.p.) and isoflurane (1.2-2% in 100% O_2_) at the time of surgery. A custom titanium headpost was cemented to the skull with Metabond (Parkell) and a 5 mm craniotomy (coordinates from lambda: 3.10mm lateral, 1.64mm anterior) was fit with a custom-made glass window (an 8-mm coverslip bonded to two 5-mm coverslips (Warner no. 1) with refractive index-matched adhesive (Norland no. 71)) using Metabond. After surgery, animals were given buprenorphine (0.05 mg/kg) and cefazolin (50 mg/kg) for 48 hours.

Mice were allowed to recover for at least 7 days post-surgery before beginning habituation to head restraint. Habituation to head restraint increased in duration from 15 min to >2 h over 1-2 weeks. During habituation and imaging sessions, mice were head restrained while allowed to freely run on a circular disc (InnoWheel, VWR). After habituation and retinotopic mapping, wild-type mice used for axon imaging experiments (n = 5) were injected with virus to express GCaMP6s selectively in V1 neurons. Dexamethasone was administered at least 2 h before surgery and animals were anesthetized with isoflurane (1.2-2% in 100% O_2_). The cranial window was removed and a glass micropipette was filled with virus (AAV1.Syn.GCaMP6s.WPRE.SV40, 6.18×10^13^ (UPenn)), mounted on a Hamilton syringe, and lowered into the brain. 100 nL of virus was injected at 250 and 450 µm below the pia (100 nL/min); the pipette was left in the brain for an additional 10 minutes to allow the virus to infuse into the tissue. Following injection, a new coverslip was sealed in place. Imaging experiments were conducted 6-24 weeks following injection to allow for sufficient expression.

### Visual Stimulation

Visual stimuli were presented on a 144-Hz LCD monitor (Asus) calibrated with an i1 Display Pro (X-rite). The monitor was positioned 21 cm from the contralateral eye. Presentation of drifting, vertical, circularly-apertured, sine-wave gratings alternated with periods of uniform mean luminance (60 cd/m^2^) and was controlled with MWorks software (http://mworks-project.org).

For widefield imaging of retinotopy to identify V1 and the HVAs, 30° diameter sinusoidal gratings (spatial frequency (SF): 0.1 cycles per degree (cpd); temporal frequency (TF): 2 cycles per second (Hz); vertical and drifting right-ward) were presented in a 3×3 grid with 15-20° spacing (Figure 1A) for 5 s with 5 s inter-trial interval (ITI) for GCaMP imaging or 10 s with a 10s ITI for autofluorescence imaging.

For two-photon (2P) imaging experiments, the SF and TF of the stimuli were tailored to the tuning of the neurons in that area: V1: 0.1 cpd and 2 Hz; AL and LM: 0.04 cpd and 6 Hz; PM: 0.16 cpd and 1 Hz (Andermann et al., 2011). When measuring the fine spatial receptive fields of neurons, we presented vertical drifting gratings in a 7×7 evenly spaced grid (Figure 1C) and varied the diameter and spacing of the stimuli according to area: V1: 10° diameter/5° spacing; LM, AL and PM: 20° diameter/10° spacing. When imaging axons in the HVAs, V1 stimulus parameters were used independent of target area. In each experiment, the position that most effectively activated the imaged population was chosen for the size-tuning experiment. Effort was made to have the stimulus be centered on the monitor; however, in some cases, parts of the larger stimuli were cut off on the edges. For most size-tuning experiments we examined 8 sizes and 4 contrasts (log-spaced from 0.1 to 0.8); in a subset of experiments we used (when imaging V1 axons in the HVAs) only the highest contrast (0.8). All stimuli during 2P imaging were presented for 1 s with a 3 s ITI, and all stimulus conditions were randomly interleaved and repeated at least 10 times.

### Widefield and two-photon imaging

Retinotopic maps of V1 and the HVAs (Figure 1B) were generated using widefield imaging of either intrinsic autofluorescence or GCaMP signals. The brain was illuminated with blue light (473 nm LED (Thorlabs) with a 462 ± 15 nm bandpass filter (Edmund Optics)), and emitted light was measured through a 500 nm longpass (for autofluorescence) or a 520 ± 18 nm bandpass filter (for GCaMP). Images were collected using a CCD camera (Rolera EMC-2, Qimaging) at 2 Hz through a 5x air immersion objective (0.14 numerical aperture (NA), Mitutoyo), using Micromanager acquisition software (NIH). Images were analyzed in ImageJ (NIH) to measure changes in fluorescence (dF/F; with F being the average of all frames) to identify primary visual cortex (V1) and the higher visual areas. Vascular landmarks were used to identify targeted sites for 2P imaging or viral injections.

2P images of neurons and axons in the visual cortex were collected using a microscope controlled by Scanbox software (Neurolabware). Excitation light (920 nm) from a Mai Tai eHP DeepSee laser (Newport) was directed into a modulator (Conoptics) and raster scanned onto the brain with a resonant galvanometer (8 kHz, Cambridge Technology) through a 16X (0.8 NA, Nikon) water immersion lens. Average power at the surface of the brain was 30-50 mW. Frames were collected at 15.5 Hz for a field of view (FOV) of approximately 605 × 342 μm for cell bodies and 500 × 300 μm for axons. Emitted photons were directed through a green filter (510 ± 42 nm band filter (Semrock)) onto GaAsP photomultipliers (H10770B-40, Hamamatsu). For cell bodies, images were captured at a plane 250 ± 50 μm below the pia (range 200-300 μm; n = 9 mice, 52 FOV (V1: 18; LM: 9; AL: 10; PM: 15)); for axons, images were 150 μm below the pia (LM: 4 mice, 4 FOV; AL: 3 mice, 3 FOV; PM: 4 mice, 7 FOV). Frame signals from the scan mirrors were used to trigger visual stimulus presentation for reliable alignment with collection.

## Data Analysis

All 2P imaging data was analyzed using custom code written in MATLAB (Mathworks).

### Registration and segmentation

Image stacks from each imaging session were registered for x-y motion to the same stable reference image selected out of several 500-frame-average images, using Fourier domain subpixel 2D rigid body registration.

Cell bodies were manually segmented from a filtered (3×3 pixel median filter) image of the average dF/F during the 1 s of stimulus presentation (where F is the average of the the last 1 second of the ITI) for each stimulus presented during the fine retinotopy experiment. Fluorescence time courses were generated by averaging all pixels in a cell mask. Neuropil signals were removed by first selecting a shell around each neuron (excluding neighboring neurons), estimating the neuropil scaling factor (by maximizing the skew of the resulting subtraction), and removing this component from each cell’s time course. Visually evoked responses were measured as the average dF/F during a 1 s window around the peak response (exact window was selected separately for each experiment to account for variability in response latencies and indicator kinetics) where F is the average of the 1 s preceding the stimulus.

Axons were automatically segmented from the filtered, average dF/F images acquired during the fine retinotopy. Single pixels were identified as the center of an axonal bouton if they met the following criteria: 1) was the brightest pixel of the nine neighboring pixels, 2) had a dF/F of at least 0.05, and 3) was significantly responsive to at least two stimulus positions. Masks for each bouton included the single pixel plus the nine surrounding pixels. Neighboring boutons could be no less than 5 pixels from center to center, so that there were no pixels included in more than one bouton. The same approach as for cell bodies was used to extract time courses and measure single trial responses, except no neuropil subtraction was performed on the boutons. In the case that there were one or two retrogradely labeled cell bodies in the HVAs, the area around the cell body was blanked for segmentation; in the case that there were more than two cell bodies, the experiment was discarded.

Following segmentation, the same analyses were performed on both cells and boutons unless otherwise stated. For both cells and boutons, the masks found in the retinotopy experiment were applied to the images collected during the size-tuning experiment.

### Fitting Spatial Receptive Fields (RFs)

The retinotopic organization of single cells and boutons was assessed by measuring the average dF/F response to each of 49 stimulus positions (7×7 grid). These data were fit by least-squares regression with a 2-dimensional Gaussian curve:

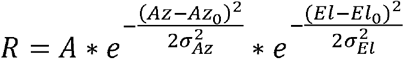

where R is dF/F response, Az is stimulus azimuth, El is stimulus elevation, A is Gaussian amplitude, Az_0_ is RF center in azimuth, El_0_ is RF center in elevation, **σ**_Az_ is standard deviation of RF in azimuth, **σ**_El_ is standard deviation of RF in elevation. Receptive field size was calculated as the geometric mean of the half-width at half-maximum (HWHM) in each dimension of the Gaussian fit.

Quality and consistency of fit were assessed by resampling trials with replacement 500 times. Only cells with 95% of the RF center estimates within one step size in each dimension from the RF center fit using all of the data were included in further analysis. Additionally, cells for which the RF center estimates were within 1 degree of the edge of the grid were discarded. In the case that there were more than 2000 boutons in a field of view that were not on the edge, only boutons for which the r^2^ of the original fit was greater than 0.5 were assessed with the resampling analysis. For cell bodies, 1877/7107 in V1, 832/2745 in LM, 1003/2613 in AL, and 711/2902 in PM passed our criteria for inclusion; for V1 boutons, 3243/12581 in LM, 3815/12551 in AL, and 5083/19349 in PM passed our criteria for inclusion.

### Fitting Size-Tuning Curves

Neurons that were well-fit for spatial receptive field were subsequently fit for size. There are two major modes of responses to stimuli of increasing size that are typically observed in the visual cortex (Sceniak et al., 1999): one of a saturating response up to some size and remaining active at larger sizes (non-suppressed cell), and one of a peak response and declining response at larger sizes (suppressed cell). Thus, each cell was fit with both a saturating, single sigmoid (SS model):

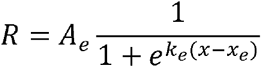

or with a suppressed, difference of sigmoids (DOS model):

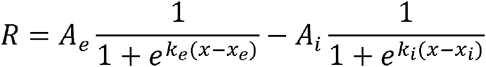

where R is dF/F response, x is stimulus size, A_e_ is excitatory sigmoid amplitude, k_e_ is excitatory sigmoid steepness, x_e_ is excitatory sigmoid center, A_i_ is inhibitory sigmoid amplitude, k_i_ is inhibitory sigmoid steepness, and x_i_ is inhibitory sigmoid center. Some parameters were constrained when fitting these models. In both models, A_e_>0 and x_e_>0 ensured a positive excitatory response centered at a size above zero. In the DS model, k_e_>k_i_ and x_e_<x_i_ ensured the second inhibitory sigmoid was less steep than and centered at a larger size than the excitatory sigmoid, to represent the larger size and spatial offset of the surround field. Initial guesses for sigmoid amplitudes and center were set based on the amplitude and size of the peak response across all sizes.

For each cell, size-tuning curves at each contrast condition were individually fit with both models using a least-squares regression method with an additional smoothness penalty to prevent overfitting of data. A sequential F-test was used to assess if the DOS fit was significantly improved from the SS fit. If the fit passed the F-test, the cell was designated suppressed (or DOS); otherwise, the cell was non-suppressed (or SS). For the SS model, preferred size was defined as the size at which 90% saturation is reached; for the DS model, preferred size is the size that generates the peak response. In the SS model, suppression index (SI) is 0; in the DOS model, SI was defined as:

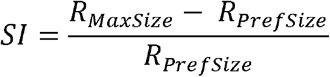

Where R_MaxSize_ is the amplitude of the response to the largest stimulus, and R_PrefSize_ is the amplitude of the response at the preferred size.

Quality and consistency of fits were again assessed by bootstrapping over 500 shuffles in the highest contrast condition, examining that the preferred size estimates remain within one octave (ratio of 2) in either direction from the unshuffled fit, and that at least 50% of fits were designated by sequential F-test as the same model as the unshuffled fit, otherwise discarding that cell from analysis. Additionally, boutons that were not significantly modulated across contrasts (according to a one-way ANOVA) were discarded from analysis. For cell bodies, 1136/1877 in V1, 446/832 in LM, 486/1003 in AL, and 351/711 in PM passed our criteria for inclusion; for V1 boutons, 2244/3243 in LM, 1869/3815 in AL, and 1514/5083 in PM passed our criteria for inclusion. Cells were also filtered from analysis based on the distance of the cell’s receptive field center (as measured from the fit) to the center of the visual stimulus position used for the size-tuning experiment. Cutoffs for the maximum distance were chosen separately for each area based on average receptive field size: 10° for V1 (and V1 axons in HVAs), 15° for LM and AL, and 20° for PM. For cell bodies, 831/1136 in V1, 202/446 in LM, 260/486 in AL, and 236/351 in PM fell within the cutoff ranges; for V1 boutons, 2022/2244 in LM, 1702/1869 in AL, and 1251/1514 in PM fell within the cutoff ranges. Additional comparisons were performed using a matched cutoff of 15° for cell bodies in all areas: 1092/1136 in V1, 202/446 in LM, 260/486 in AL, and 169/351 in PM fell within the matched cutoff range.

For analyses of size-tuning parameters across all four contrast conditions, cell bodies were selected based of the goodness-of-fit at the lowest contrast condition, in order to eliminate cells with noisy size-tuning curves at low contrasts. This was achieved by requiring r^2^ of the designated size-tuning model fit at the lowest contrast to be greater than 0.2. For cell bodies, 233/831 in V1, 87/202 in LM, 81/260 in AL, and 113/236 in PM passed our criteria for inclusion.

### Fitting Contrast Response Functions

Contrast responses were extracted from the size-tuning curve fits in each contrast condition at the preferred size of the highest contrast condition. These data were fit with a Naka-Rushton hyperbolic function:

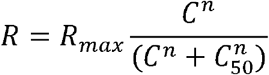

where R is dF/F response, C is stimulus contrast, n is exponent of power function (constrained >0), and C_50_ is contrast of half-max response.

### Experimental Design and Statistical Analysis

Data were tested for normality using a Lilliefors test. None of the receptive field parameters were normally distributed; thus we used the Wilcoxon rank sum and Kruskal-Wallis tests (with post-hoc Tukey tests) for two- and multiple-sample statistics. However, in the cases where we were interested in the relationship between two variables, we used a two-way ANOVA which has been shown to be robust to non-normality (Driscoll, 1996). Sample sizes were not predetermined by statistical methods, but our sample sizes of the neurons and behavior animals are similar to other studies. The numbers of cells, animals or experiments are provided in the corresponding text, figures and figure legends. All error values in the text are standard deviation unless otherwise specified. Data collection and analysis were not performed blind to experimental conditions, but all visual presentation conditions were randomized.

### Data and code availability

All relevant data and code will be made available upon reasonable request.

## Acknowledgements

We thank E. Burke for surgical assistance and members of the Hull and Glickfeld labs for helpful discussions. We thank Drs. Court Hull and Kevin Franks for comments on the manuscript. This work was supported by grants from the Pew Biomedical Trusts (L.L.G.), the Alfred P. Sloan Foundation (L.L.G), and the NIH: Director’s New Innovator Award (DP2-EY025439, L.L.G.) and Ruth L. Kirschstein Pre-Doctoral Fellowship (F31-EY028018-2, A.M.W.).

## Author contributions

K.A.M, A.M.W., V.M., and L.L.G. designed the experiments. V.M. performed pilot experiments and K.A.M. and A.M.W. collected the data. K.A.M. and L.L.G. analyzed the data. K.A.M, A.M.W. and L.L.G. wrote the manuscript.

